# Natural Genetic Variation Modifies Gene Expression Dynamics at the Protein Level During Pheromone Response in Saccharomyces cerevisiae

**DOI:** 10.1101/090480

**Authors:** Daniel A. Pollard, Ciara K. Asamoto, Homa Rahnamoun, Austin S. Abendroth, Suzanne R. Lee, Scott A. Rifkin

## Abstract

Heritable variation in gene expression patterns plays a fundamental role in trait variation and evolution, making understanding the mechanisms by which genetic variation acts on gene expression patterns a major goal for biology. Both theoretical and empirical work have largely focused on variation in steady-state mRNA levels and mRNA synthesis rates, particularly of protein-coding genes. Yet in order for this variation to affect higher order traits it must lead to differences at the protein level. Variation in protein-specific processes including protein synthesis rates and protein decay rates could amplify, mask, or even reverse effects transmitted from the transcript level, but the extent to which this happens is unclear. Moreover, mechanisms that underlie protein expression variation under dynamic conditions have not been examined. To address this challenge, we analyzed how mRNA and protein expression dynamics covary between two strains of *Saccharomyces cerevisiae* during mating pheromone response. Although divergent *steady-state* mRNA expression levels explained divergent *steady-state* protein levels for four out of five genes in our study, the same was true for only one out of five genes for expression *dynamics*. By integrating decay rate and allele-specific protein expression analyses, we resolved that expression divergence for Fig1p was caused by genetic variation acting in *trans* on protein synthesis rate, expression divergence for Ina1p was caused by *cis*-by-*trans* epistatic effects on transcript level and protein synthesis rate, and expression divergence for Fus3p and Tos6p were caused by divergence in protein synthesis rates. Our study demonstrates that steady-state analysis of gene expression is insufficient to understand the impact of genetic variation on gene expression variation. An integrated and dynamic approach to gene expression analysis - comparing mRNA levels, protein levels, protein decay rates, and allele-specific protein expression - allows for a detailed analysis of the genetic mechanisms underlying protein expression divergences.

## INTRODUCTION

Heritable variation in gene expression patterns plays a fundamental role in trait variation and evolution. From human lactase persistence (1) and disease susceptibility (2,3) to adaptive skeletal variation in stickleback fish (4), genetic variation commonly acts on traits by altering when, where, and how much genes are expressed (5,6). Understanding the mechanisms by which genetic variation acts on gene expression patterns is therefore a major goal for the biological sciences.

The expression level of a protein in a cell depends on the rates of four general processes: mRNA synthesis, mRNA decay, protein synthesis, and protein decay. Heritable genetic variation can in theory affect any of these complex molecular processes (Fig 1), resulting in differences in how cells control the expression of their genes and ultimately in how they respond to their environment. However, due in large part to the relative maturity of high-throughput RNA measurement technologies, empirical and theoretical work has focused on mechanisms that act on steady-state mRNA levels or mRNA synthesis rates (6-10). Whether variation in protein synthesis rate and protein decay rate contribute to gene expression variation and, in particular, *dynamic* (non-steady-state) gene expression, remains enigmatic.

**Fig 1.**
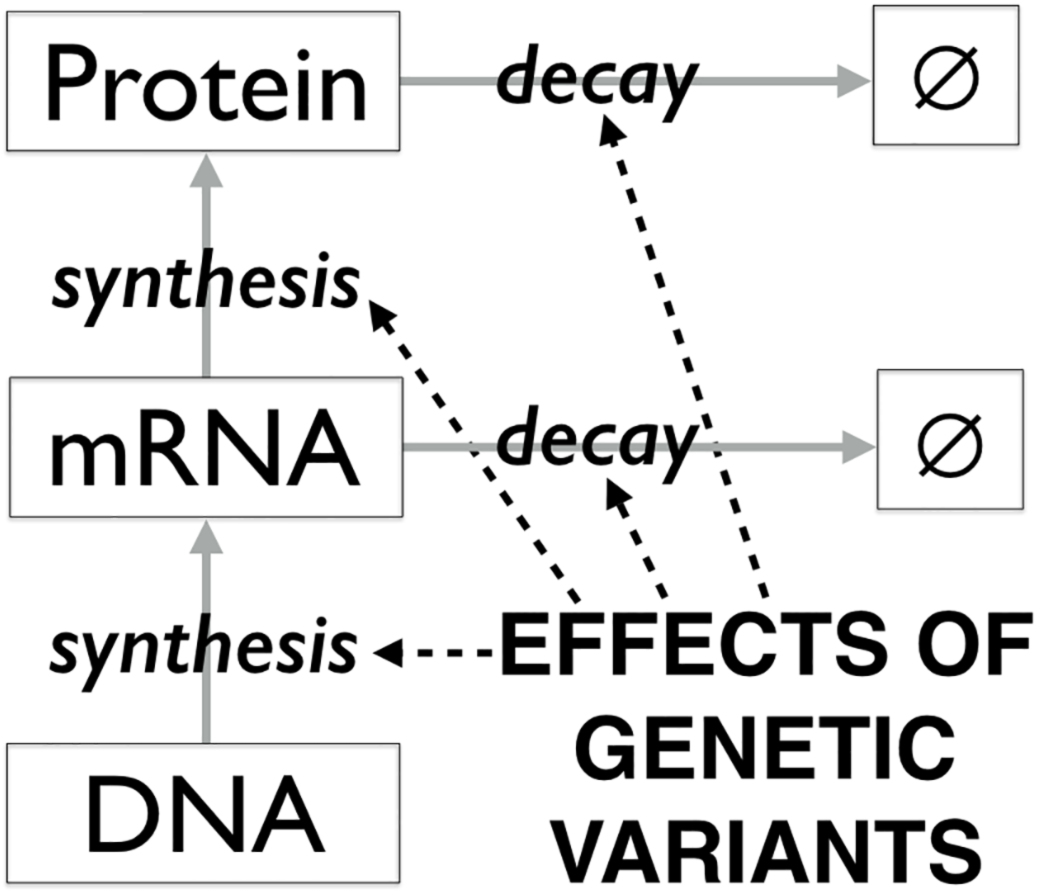
Mechanisms of gene expression variation. Genetic variants can act on gene expression by affecting the rates of mRNA synthesis, mRNA decay, protein synthesis, and protein decay.

The connection between heritable variation in mRNA levels and protein levels is an active area of investigation but has overwhelmingly focused on steady-state conditions. Initial proteomic surveys of natural variation in steady-state protein levels have found that mRNA variation is not well correlated with protein variation (11-18). A mass spectroscopy study of genetic variation in yeast underlying steady-state protein levels for highly expressed genes found that a minority of variants were linked to mRNA variation (12). Although differences in the statistical power of these methods makes direct comparisons challenging, other studies of the same yeast strains using fluorescent protein levels instead of mass spectroscopy found that only half of genetic variation affecting protein levels also affects mRNA levels (19,20), suggesting that this disconnect between mRNA and protein variation is not simply the result of technical limitations of mass spectrometry compared to microarrays or other RNA measurement technologies. Recent analyses of translation rate variation using ribosome occupancy under steady-state conditions have revealed that genetic variation can act at the protein level to either buffer (21,22) or reinforce (23) variation at the mRNA level.

Although these studies establish that steady-state mRNA variation is not sufficient to explain steady-state protein expression, it is not clear how mRNA expression variation, protein synthesis rate variation, and protein decay rate variation *combine* to produce protein expression variation. Furthermore, because these studies only focused on steady-state conditions, we do not know whether the relationship between mRNA and protein variation is different during dynamic processes such as development, disease progression, or response to environmental fluctuations.

In this paper, we analyzed how mRNA and protein expression dynamics vary between two strains of *Saccharomyces cerevisiae* and characterized cases of genetic variation acting at the protein level. We focused on five genes with known transcriptional responses to alpha-factor mating pheromone in MATa haploids. *FIG1*(YBR040W)(24), *FUS3*(YBL016W)(25), and *GPH1*(YPR160W)(26) are transcriptionally activated by alpha-factor while *INA1*(YLR413W)(27) and *TOS6*(YNL300W)(28) are repressed at the transcript level by alpha-factor (29). We compared expression dynamics between the laboratory strain S288c and the clinical isolate YJM145 (also referred to as YJM789 (30)) for which differences in their MAP kinase pathway and mRNA expression responses to alpha-factor have been characterized (31,32). We found that differences in mRNA expression dynamics were unable to explain differences in protein expression dynamics for *FIG1*, *FUS3*, *INA1*, and *TOS6*. We then characterized the relative influence of protein synthesis (33) and protein decay (34) rate variation and whether these rates vary due to *cis-acting* (allele-specific) or *trans-acting* (non-allele-specific) genetic variants or a combination of both(35).

## RESULTS

### Divergence in mRNA and protein expression dynamics

We initially sought to (a) evaluate how well divergence in mRNA expression dynamics can explain divergence in protein expression dynamics and (b) identify genes with evidence for divergence in protein synthesis rate or protein decay rate. We measured the expression level of mRNA and protein for the genes *FIG1*, *FUS3*, *GPH1*, *INA1*, and *TOS6* during a time-course of alpha-factor mating pheromone response in MATa haploids of the yeast strains S288c and YJM145 (Fig 2A-J). Cells were collected from a mid-log culture of each strain throughout an eight hour exposure to alpha-factor (50nM final). We measured relative mRNA concentration using RT-qPCR (Fig 2A-E) and relative protein concentration using fluorescence microscopy (Fig 2F-J) followed by Western Blots for confirmation (S1 Figure). Each time-course experiment for each gene was performed in triplicate to evaluate the repeatability and significance of mRNA and protein expression divergences between strains. In addition to the ratio of mRNA and protein expression between strains (Fig 3A-J), we isolated the divergence in protein expression from the divergence in mRNA expression by calculating the divergence in protein units per mRNA unit between strains (Fig 3K-O).

**Fig 2.**
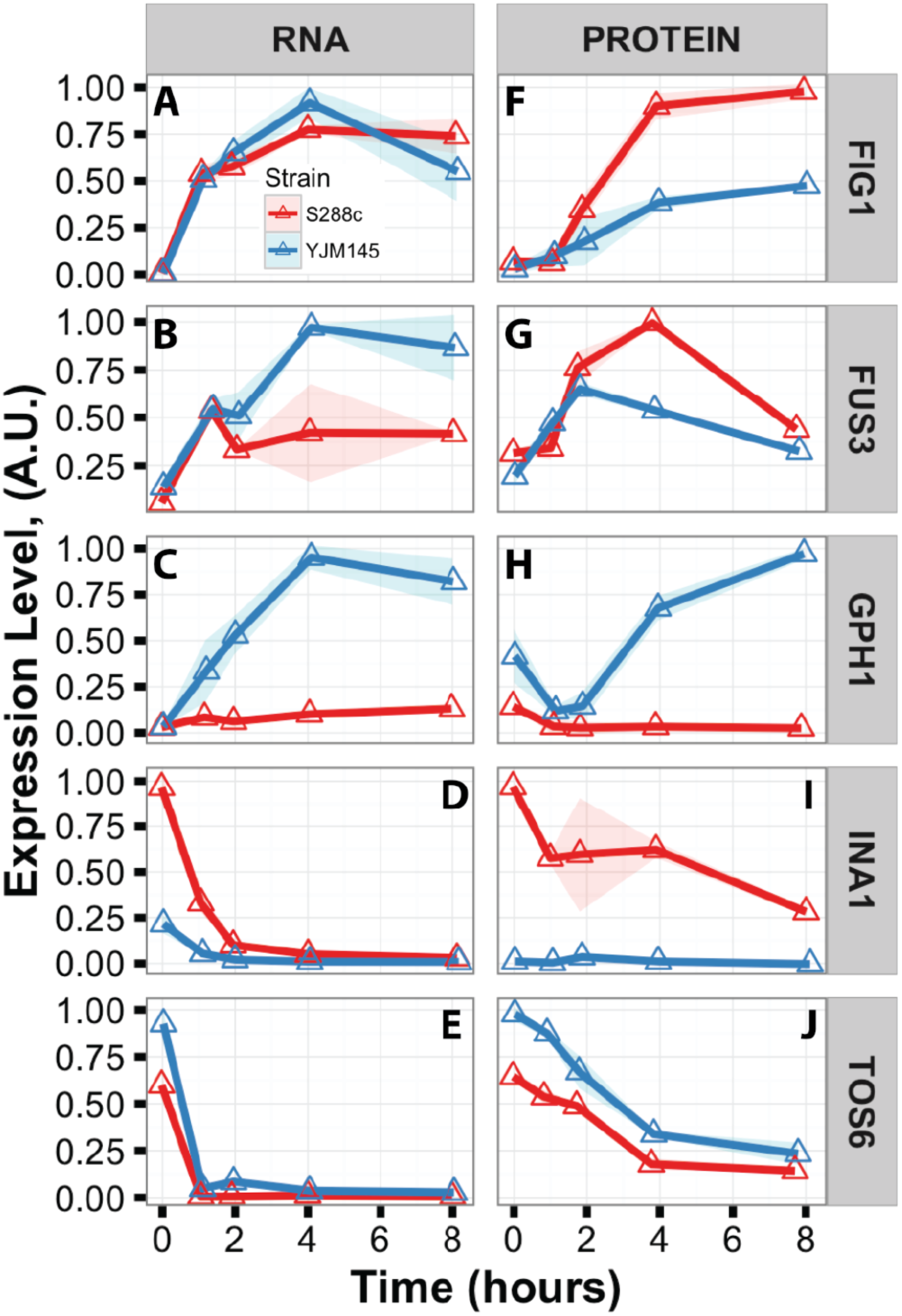
Example mRNA and protein expression responses to pheromone in strains S288c and YJM145. (A-J) Example RNA and protein expression time-courses in S288c (red triangles, lines, and shadows) and YJM145 (blue triangles, lines, and shadows) following addition of pheromone at time 0hr. Triangles are mean expression over technical triplicates and shadows are 95% confidence intervals. All plots are rescaled to arbitrary units (a.u.) of a maximum value of one for visualization purposes.

mRNA expression dynamics significantly differed between strains across multiple time-points for the genes *GPH1*, *INA1*, and *TOS6* (Fig 3C,D,E) while protein expression dynamics significantly differed across multiple time-points for all five genes (Fig 3F-J). For *FIG1*, mRNA expression was very similar between strains (Fig 2A and 3A) while protein expression was significantly higher in S288c than YJM145 after an hour of pheromone treatment (Fig 2F, 3F, and Fig A,D,E,J in S1 Figure). *FIG1* protein per mRNA significantly diverged between the two strains over most of the time-course (Fig 3K), indicating that genetic variation is acting directly on protein synthesis and/or protein decay. For *FUS3*, mRNA expression trended toward lower expression in S288c (Fig 2B and 3B) while protein expression was significantly higher in S288c for most time-points by microscopy and trended higher by Western blot (Fig 2G, 3G, and Fig B,F,G,J in S1 Figure). Furthermore, FUS3 protein per mRNA was significantly higher in S288c for most time-points (Fig 2G, 3G,L, and Fig B,F,G,J in S1 Figure), all together indicating another likely case of variation in protein synthesis and/or protein decay rates between strains. Although *GPH1* had dramatically different expression dynamics in the two strains, protein expression divergence mirrored mRNA expression divergence (Fig 2C,H and 3C,H,M), indicating that genetic variation affecting *GPH1* acts predominantly at the transcript level. For *INA1*, both mRNA and protein expression were significantly higher in S288c than YJM145 for most time-points (Fig 2D,I, 3D,I, and Fig C,H,I,J in S1 Figure). However, at the time point prior to adding pheromone, protein per mRNA was significantly lower in YJM145 (Fig 3N), due to the modest quantity of mRNA produced by YJM145 resulting in protein levels at the threshold of detection, whether assayed by fluorescence microscopy or Western blot (Fig 2D,I and Fig C,H,I,J in S1 Figure). Finally, both *TOS6* mRNA and protein had lower expression in S288c relative to YJM145 (Fig 2E,J and 3E,J), but the magnitude of the mRNA expression divergence was greater than that of the protein expression divergence (Fig 3E,J,O), again suggesting that genetic variation is acting at the protein level.

**Fig 3.**
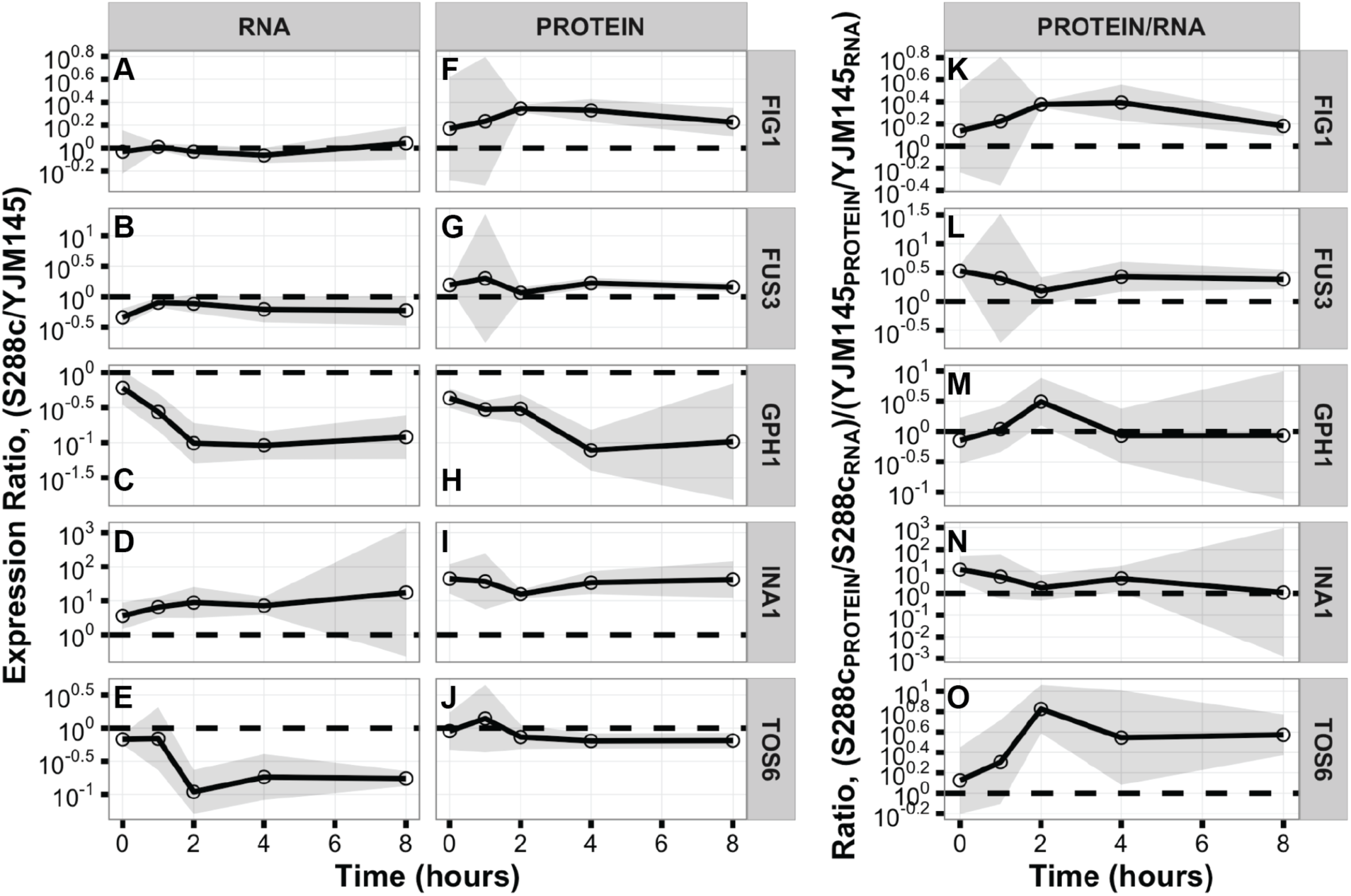
Comparison of average expression responses to pheromone between strains S288c and YJM145. (A-J) Ratio of mRNA or protein expression in S288c over YJM145. (K-O) Ratio of protein per RNA in S288c over YJM145. Circles are mean ratio over biological triplicates and shadows are 95% confidence intervals. Dashed horizontal line is ratio of 1. Y-axes are log_10_ scale.

Our analysis of mRNA and protein expression dynamics for five genes suggests four clear examples of genetic variation that specifically acts on protein synthesis and/or protein decay rates. We next sought to further characterize the protein expression divergences in *FIG1*, *FUS3*, *INA1*, and *TOS6*.

### Divergence in protein synthesis versus protein decay rates

Our next goal was to evaluate the relative role of protein synthesis rate variation and protein decay rate variation for the proteins Fig1p, Fus3p, Ina1p, and Tos6p. We applied the protein synthesis inhibitor cycloheximide (20µg/ml final) to cultures of each strain to block protein synthesis and then measured relative protein concentration over two hours using fluorescence microscopy. We added cycloheximide in the absence of pheromone for Ina1p decay assays and three hours into pheromone treatment for Fig1p, Fus3p, and Tos6p decay assays, time-points when protein per mRNA for each gene was highly diverged (Fig 3).

None of the four proteins had significant differences in protein decay rates between strains (Fig 4). Fig1p, Ina1p, and Tos6p protein molecules appear very stable in both strains, with decay rates close to zero (Fig 4A,C,D,E). In contrast, Fus3p protein molecules decay rapidly, with similar rates in both strains (Fig 4B).

**Fig 4.**
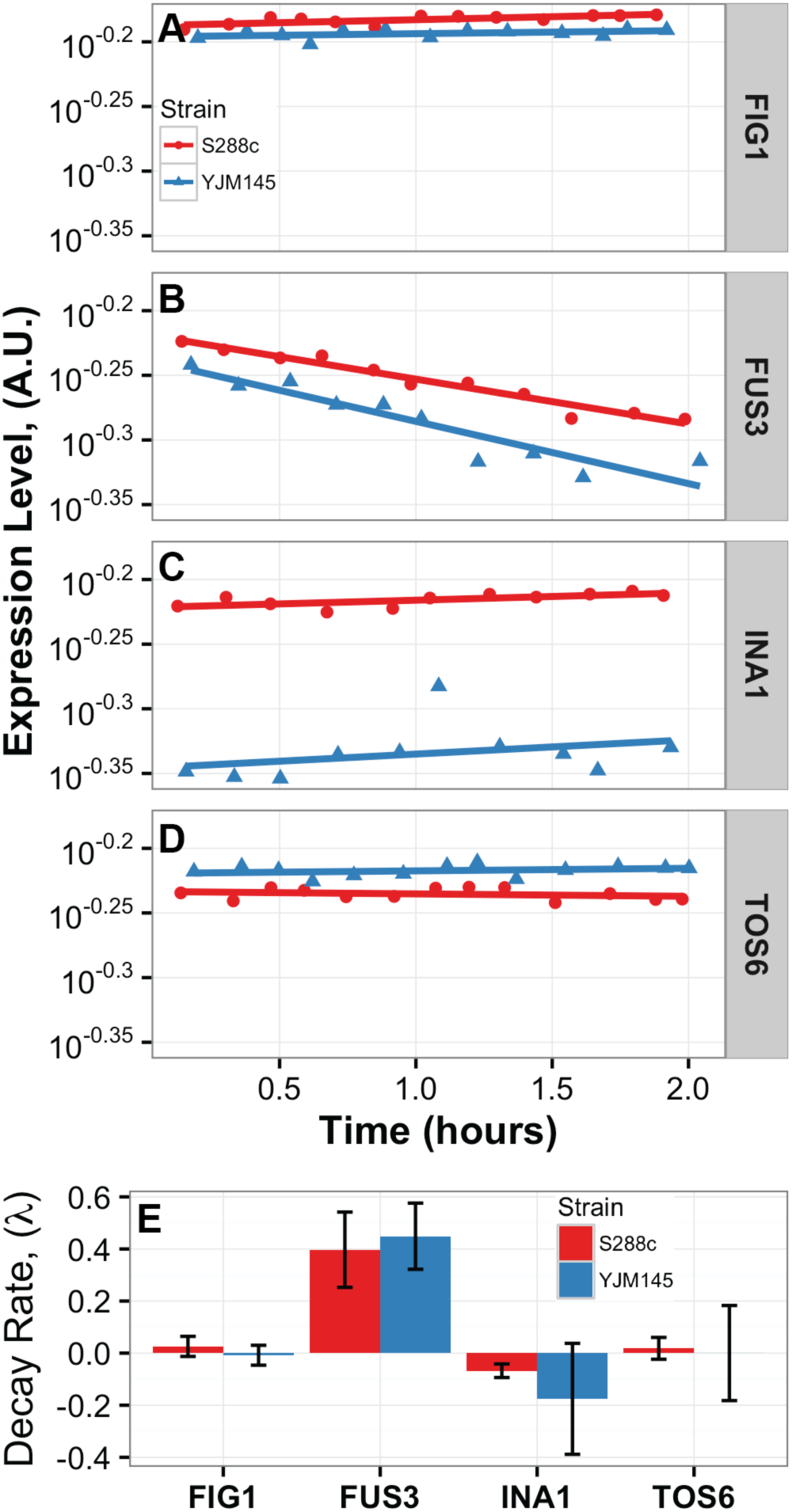
Relative protein decay rate between strains S288c and YJM145. (A-D) Example protein decay time-courses in S288c (red circles) and YJM145 (blue triangles). Pheromone added at time −3hr for Fig1p (A), Fus3p (B), and Tos6p (D). Cycloheximide added at time 0hr. Circles and triangles are log_10_ of expression. Lines are linear regression on log_10_ of expression through time. (E) Decay rate parameter lambda estimated from the negative slope of the linear regression line fit to log_10_ of expression through time. Bar heights are mean lambda over biological triplicates and error bars are 95% confidence intervals.

The lack of evidence for significant differences in protein decay rates for Fig1p, Fus3p, Ina1p, and Tos6p indicate that genetic variation acts on protein synthesis rates for these genes. *Cis-acting versus trans-acting genetic variation.*

Our final goal was to determine whether the genetic variation affecting protein expression patterns is *cis*-acting or *trans*-acting. We focused on *FIG1* and *INA1*. We devised an allele-specific protein expression assay to detect *cis*-acting genetic variation. For each gene we added a second copy of the gene fused to a different fluorescent protein in a different genomic location. This gave us four derived strains for each gene - two with heterologous alleles in the two locations used to measure allelic effects on expression and two with identical alleles used to control for fluorophore intensity and genomic location. For *FIG1* we compared protein expression of each allele during a time-course of pheromone response using fluorescence microscopy. We compared allelic expression in *INA1* cultures without pheromone, the time-point when divergence in protein per mRNA was greatest.

*FIG1* showed no evidence of allele-specific protein expression. The ratio of expression levels of the two strain alleles was close to one in both strain backgrounds (Fig 5A), consistent with *trans*-acting variation. *INA1* had a small but significant difference in the protein expression of alleles in the S288c strain background (Fig 5B and S2 Figure) while alleles had low and indistinguishable expression in the YJM145 strain background (Fig 5B and S2 Figure). Thus *INA1*’s low protein expression in YJM145 appears to be due to a combination of genetic variation acting in *cis* and in *trans*.

**Fig 5.**
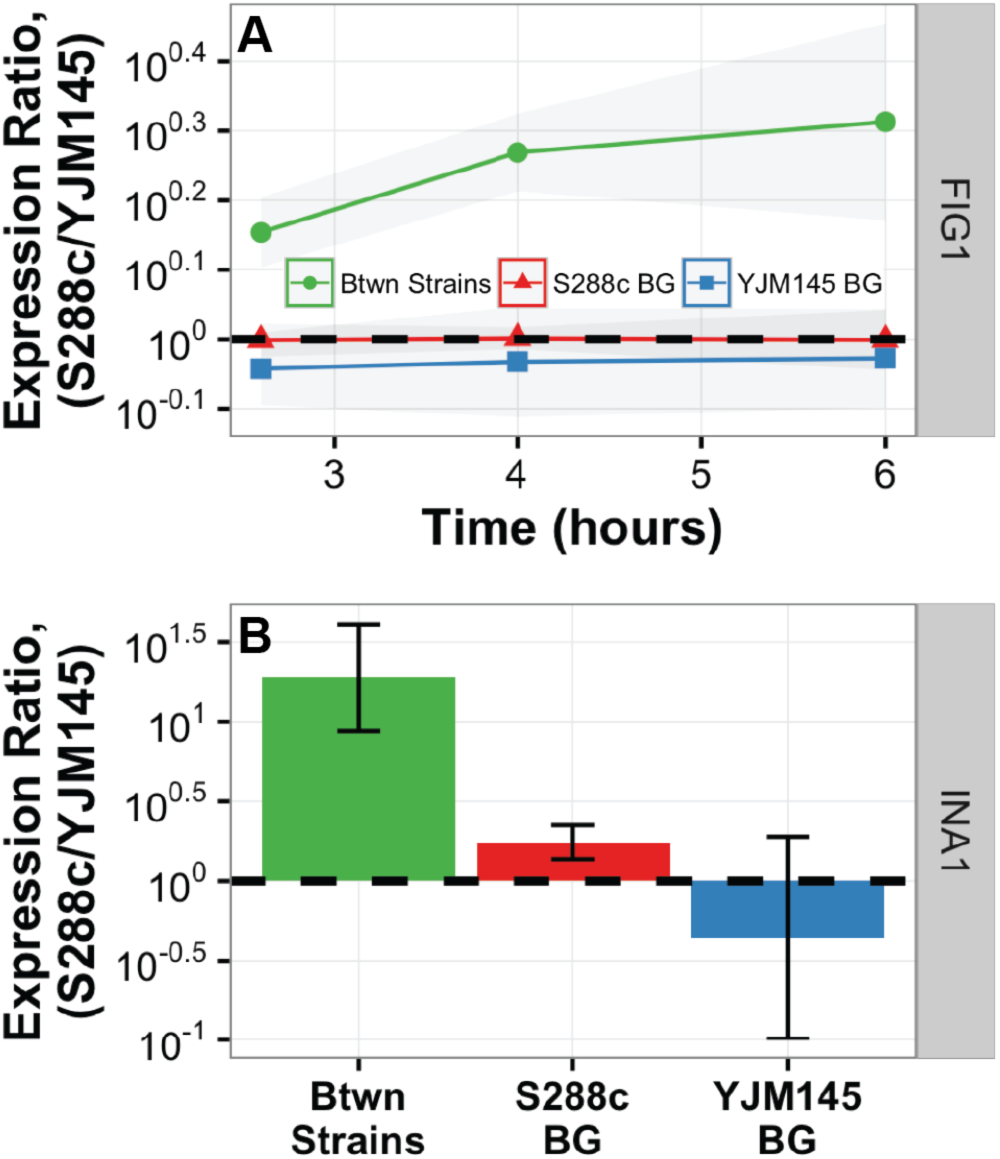
Allele-specific protein expression. (A) Ratio of S288c to YJM145 Fig1p protein allele expression within and between strain backgrounds following addition of pheromone at time 0hr. Symbols are mean ratio over biological triplicates and shadows are 95% confidence intervals. Horizontal line is ratio of 1. Y-axis is log_10_ scale. (B) Ratio of S288c to YJM145 Ina1p protein allele expression within and between strain backgrounds during steady-state growth. Bar heights are mean ratio over biological triplicates and error bars are 95% confidence intervals. Horizontal line is ratio of 1. Y-axis is log_10_ scale.

## DISCUSSION

The central question driving this work was: how well does natural variation in mRNA expression explain natural variation in protein expression under dynamic conditions. All five genes in our study had significant differences in protein expression responses to pheromone between strains (Fig 2 and 3), but *GPH1* was the only gene for which the variation in mRNA expression largely explained the variation in protein expression. The other four genes exhibited distinct patterns of discordance in expression divergence between mRNA and protein. We observed one case of conserved mRNA and divergent protein (*FIG1*), two cases of opposing (compensating or buffering) variation at the mRNA and protein levels (*FUS3* and *TOS6*), and one case of reinforcing variation where a quantitative divergence at the mRNA level turned into nearly a complete loss of expression in one strain at the protein level (*INA1*). From this small study of pheromone responsive genes, it is clear that the patterns of divergence at the mRNA and protein levels can be diverse. It is also clear that variation in mRNA expression dynamics is insufficient to explain protein expression dynamics for most genes (in our case 80%).

Our goal in this small study was not to provide an estimate of the global fraction of genes with genetic variation specifically targeted to protein dynamics. Instead, we wanted to explore the types of concordance and discordance between variation in mRNA and protein trajectories. To our surprise we found a variety of patterns within a handful of genes. We selected these genes because they were known to have dynamic mRNA expression patterns (29), which was important given our focus on dynamics. Previous work had also shown that *FIG1*, *FUS3*, and *GPH1* were differentially induced at the mRNA level between S288c and YJM145 30 minutes into a 5µM pheromone treatment and that *INA1* and *TOS6* were not (32). Nothing was known about any differences in protein levels. Moreover, except for *GPH1*, our measurements of mRNA divergence were inconsistent with the single time-point study of Zheng et al.(32), perhaps because of differences in strain background, pheromone dose, or other experimental factors. With respect to our central question, we had no prior information for any of the genes. Consequently, we have no reason to believe these five genes are somehow unusual, and we suspect that a large fraction of regulated genes in the genome will exhibit divergence in protein expression dynamics that cannot be explained solely by transcript level divergence.

Our first result implied a role for genetic variation acting on protein expression through a mechanism that does not impact mRNA expression levels. Processes that add or subtract protein from the cell given a pool of mRNA are protein synthesis and protein decay. Mechanisms for the protein synthesis rate variation described here could act as early as translation initiation (33) and as late as translation termination as our Western blots and fluorescence assays require a mature fluorophore for detection. Though we noted some discrepancy in absolute variation values between our Western blotting and fluorescence microscopy data (S1 Figure) - which could be the result of variation in protein folding or simply methodological biases such as solubility of membrane-associated proteins (Fig1p and Ina1p) - the trends in variation remained the same. Protein decay is inclusive of passive decay as well as regulated decay (34).

Our second goal was to categorize the mechanistic basis of variation in protein expression dynamics as occurring at the level of either protein decay or protein synthesis rate or a mixture of both. Our decay rate assay does not control for the effect cycloheximide has on the protein decay machinery itself and so is not an absolute measure of protein decay rate but a relative one between the strains (see Methods). For Fig1p, Ina1p, and Tos6p, decay rates were very low and not significantly different between strains, indicating divergence in protein synthesis rate is the more likely cause of divergence in expression of these proteins. However, for Fus3p, decay rates were high - and therefore could have contributed to protein expression divergence - but were not significantly different between strains. Thus our integrated approach enabled us to identify four cases of divergence in protein expression dynamics that are likely explained by protein *synthesis* rate variation.

Our final goal was to classify the genetic variation underlying divergence in protein expression dynamics as allele-specific *cis*-acting or non-allele-specific *trans*-acting (35). Recent analyses of ribosome occupancy and protein levels by mass spectrometry in diploid hybrids of yeast strains and species found similar proportions of *cis-* and *trans*-acting variation underlying mRNA divergence and protein synthesis rate divergence under steady-state conditions, with higher levels of *trans*-acting variation between strains and higher levels of *cis*-acting variation between species (15,21-23). We developed a microscopy-based allele-specific protein expression assay and applied it to protein expression divergences in Fig1p and Ina1p. Consistent with the genome-wide, steady-state observations of more prevalent *trans*-acting variation within populations, we observed *trans*-acting effects for both Fig1p and Ina1p expression. *FIG1* had conserved mRNA and diverged protein expression dynamics and thus the lack of allele-specific expression in Fig1p enabled us to conclude that the divergence in Fig1p protein expression dynamics can be explained by genetic variation acting in *trans* on protein synthesis rates. The results for *INA1* imply a more complicated mechanism for the divergence of Ina1p expression. Because *INA1* had diverged mRNA and even more diverged protein expression, the divergence in protein expression is likely the result of multiple genetic variants acting at both the transcript and protein levels. Furthermore, the presence of allele-specific expression in the S288c strain background and the absence of allele-specific expression in the YJM145 strain background suggest an epistatic interaction between *cis*-acting and *trans*-acting genetic variants, such that the *cis*-acting effects of *INA1* alleles are lost in the presence of the *trans*-acting YJM145 strain background. So called *cis-trans* epistasis has been observed before at the transcript level (36), however this is the first example to our knowledge of *cis-trans* epistasis acting at both the transcript and protein levels. Thus divergence in Ina1p expression appears to result from a complex epistatic interaction of genetic variants acting in *cis* and *trans* on transcript level and protein synthesis rate.

In this study we dissected how genetic variation affects divergence in protein expression under the dynamic conditions of the pheromone response. We identified contributions from mRNA variation and protein synthesis rate variation, and for two examples further characterized the contributions of *cis*-acting and *trans*-acting genetic variants. Future studies will focus on identifying the causal polymorphisms and how they impact cellular machinery. Given that protein per mRNA levels for all genes were either conserved or higher in S288c relative to YJM145, that Fig1p, Fus3p, Ina1p, and Tos6p all had evidence of divergence in protein synthesis rates, and that both Fig1p and Ina1p had evidence of *trans*-acting genetic variation, it is possible that some or all of these genes share a causal mechanism. Recently developed methods for mapping the genomic location of genetic variation underlying protein expression levels (19,20,37) could be applied to *FIG1*, *FUS3*, and *INA1*. The compensatory nature of the genetic variation acting on *FUS3* and *TOS6* protein synthesis rates may require controlling for the genetic variation acting at the transcript level in order to observe mappable segregation of protein expression levels.

Our study focused on the causes of divergence in protein expression *dynamics* as opposed to steady-state expression levels. Had we only measured steady-state expression during log growth, we would have concluded that just one out of the five genes have mRNA variation that fails to explain protein expression variation. Although convenient for replicated laboratory studies, steady-state conditions are highly artificial. Gene expression is a dynamic process (38), and the effects of polymorphisms often act at specific times (39) or influence rates of protein synthesis or decay. Including dynamics gave us a much richer view of how phenotypic variation is generated and may benefit future studies of genetic variation.

## MATERIALS AND METHODS

### Base strains

We acquired a MATa lys5(YGL154C) gal2(YLR081W) strain of S288c called ZWY01 and a MATa gal2 strain of YJM145 (30,40) called YJM789K5 from Michael Snyder at Stanford University. Using W303 alleles of LYS5 and GAL2, we created prototrophic strains with functional galactose metabolic pathways. In both strains, we deleted the genes *URA3*(YEL021W) for use as a selection marker, *AMN1*(YBR158W) to prevent clumping and facilitate single-cell imaging (41), and *BAR1*(YIL015W) to prevent active degradation of pheromone in our experiments (42,43). The resulting strains, YSR0095 (S288c MATa lys5::W303-LYS5, gal2::W303-GAL2, ura3::loxp, amn1::loxp, bar1::loxp) and YSR0096 (YJM145 MATa gal2::W303-GAL2, ura3::loxp, amn1::loxp, bar1::loxp), were used to construct all other strains in this study.

### Citrine strains

For each of the five genes in the study, we created c-terminal translational fusions with the yellow fluorescent protein gene yECitrine and a 3x HA tag in each of the two strain backgrounds (S288c and YJM145). We constructed the plasmid pDAP5 to contain the cassette linker-yECitrine-3xHA-ADH1t-TEFp-CaURA3. The cassette was PCR amplified with a forward primer containing a 40bp overlap with the last 40bp of each gene before the stop codon and a reverse primer containing a 40bp overlap with the first 40bp following the stop codon. We transformed the PCR product into YSR0095 and YSR0096 and selected on -URA media for transformants with the cassette having replaced the stop codon for the targeted gene. Integration was verified using PCR, sequencing, and fluorescence microscopy. Using this approach we constructed ten citrine strains: YSR0115 (S228 *FIG1*::yECitrine), YSR0104 (YJM145 *FIG1*::yECitrine), YSR0097 (S228c *FUS3*::yECitrine), YSR0098 (YJM145 *FUS3*::yECitrine), YSR0135 (S288c *GPH1*::yECitrine), YSR0136 (YJM145 *GPH1*::yECitrine), YSR0143 (S288c *INA1*::yECitrine), YSR0144 (YJM145 *INA1*::yECitrine), YSR0122 (S288c *TOS6*::yECitrine), and YSR0123 (YJM145 *TOS6*::yECitrine).

### Pre-experiment strain handling

Prior to each experiment described below, we streaked out strains from frozen glycerol stocks onto solid media, incubated at 30˚ for ~24 hours, picked colonies into small overnight liquid media cultures at 30˚, and diluted to appropriate cell density based on inferred cell count from OD600.

### Liquid media

We performed all experiments using low fluorescence (-riboflavin -folic acid) synthetic complete 2% glucose media. CSM and YNB supplied by Sunrise Science.

### mRNA and protein expression time-courses

We performed each time-course experiment in triplicate over three days. We cultured and handled both the S288c and YJM145 citrine strains for a given gene simultaneously in an identical manner. To measure and control for autofluorescence levels during protein quantification, we also cultured the strains YSR0095 (S288c) and YSR0096 (YJM145) that lack the yECitrine gene simultaneously in an identical manner. We started 200ml cultures at an OD600 of ~0.07 and grew them for three hours to an OD600 of ~0.28. We added alpha-factor pheromone (Zymo) to a final concentration of 50nM to the cultures and collected cells from the cultures for RNA and protein quantification just prior to adding pheromone (0hr) and then at four time-points over the following eight hours (1hr, 2hr, 4hr, and 8hr).

### RNA sample collection and total RNA preparation

We collected 50ml of culture at the 0hr time-point, calculated the cell count from 4,269,000 cells per ml per unit OD600 for S288c and 4,785,000 cells per ml per unit OD600 for YJM145, and then collected the same number of cells for the remaining four time-points (~10^7-^10^8^). Cells were spun down, aspirated, snap frozen in liquid nitrogen, and stored at -80˚. Frozen cell pellets for an entire experiment were thawed, and ribosome depleted total RNA was isolated according to the manufacturer’s instructions for the Ambion Yeast RiboPure RNA Purification Kit (AM1926).

### cDNA preparation and RT-qPCR quantification

We produced cDNA from 2µg of our RNA samples according to the manufacturer’s instructions for the Applied Biosystems High Capacity cDNA Reverse Transcription Kit (4368814). For RT-qPCR quantification, we used primers targeting yECitrine (oDAP391: CCCATACGATGTTCCTGACTATG, oDAP392: AGCACTGAGCAGCGTAATC) and the control gene RPL7AB (oDAP369: GTCTACAAGAGAGGTTTCGGTAAG, oDAP370: CCCAAGTTGGCTTCGATGATA). To normalize for variation in total cDNA, we used the control gene RPL7AB. We tested several genes with the lowest mRNA variation during pheromone response (29) and RPL7AB gave the most robust PCR product. We ran the samples for each time-course experiment in technical triplicate on the same 96-well plate together with serial dilutions of a pooled mix covering four orders of magnitude. We used 15µl reactions with Applied Biosystems Power SYBR Green Master Mix in a BioRad CFX Connect Real-Time PCR Detection System. Using the BioRad CFX Manager we calculated SQ (Starting Quantity) values for each non-serial dilution well. We calculated expression levels for each technical replicate of each sample by dividing the yECitrine primers SQ value by the average over technical replicates of the RPL7AB primers SQ values.

### Protein sample collection

We collected 1ml of culture and concentrated or diluted the sample to ~10^6^ cells in 100µl of media based on OD600. We pre-treated wells in a 96-well glass bottom plate with 1.5mg/ml Concanavalin A. We plated samples into pre-treated wells, pelleting at 200g for 2 minutes, and then imaged immediately.

### Protein imaging and quantification

We imaged three randomly selected positions in each sample well at 60x or 63x. For each position we acquired a single in focus fluorescence image and a stack of brightfield images for automatic segmentation. In order to get the average fluorescence level within individual cells for many images, we automatically segmented the images using a version of an existing algorithm (43) modified in Matlab to handle expression patterns in our shmooing cells. For our modifications we considered many z-slices away from the focal plane in order to capture the shape of the neck and shmoo tip and we also swelled the outline of the cell three pixels in order to fully include the cell membrane where both *FIG1* and *INA1* localize. For each segmented cell in an image, we calculated mean fluorescence pixel intensity. We used fluorescence levels in our base strains YSR0095 and YSR0096 that lack the yECitrine gene, and therefore only autofluoresce, to calculate two statistics: minimum mean pixel intensity and expected autofluorescence. The minimum mean pixel intensity was used to identify segmented objects that are not cells. If the distribution of mean pixel intensities for YSR0095 or YSR0096 was monomodal then we used the 0.01 quantile of the distribution for this threshold. If the distribution was bimodal, we fit a mixture of two gaussians and used the 0.01 quantile of the gaussian with the larger mean. Similarly, we set the autofluorescence at the median of the distribution when monomodal or the mean of the gaussian with the larger mean when bimodal. Using these two statistics, we filtered out non-cells and subtracted off the autofluorescence level from our fluorescence samples. We used the mean over cells of the mean pixel intensity for each of the three image positions for downstream analysis.

### RNA and Protein time-course alignment and statistical analysis

For accurate statistical comparison of RNA and protein expression values between biological replicates performed on different days with similar but not identical time-points, we estimated the expression level at the idealized time-points of 0hr, 1hr, 2hr, 4hr, and 8hr. To do this, we constructed 243 unique paths through the technical triplicates at the five time-points, estimated the expression level at the idealized time-points on each path using linear interpolation, and then used the mean estimated value over the 243 paths as the expression level. Because the S288c and YJM145 strains were paired together for each biological replicate, we used a paired t-test in R(44), comparing the log ratio of expression between the strains across biological replicates to an expected value of zero.

### Western Blot analysis

For *FIG1* and *FUS3* strains, we added alpha-factor (50nM final) to mid-log cultures,and harvested 10^8^ cells after three hours. As *INA1* is suppressed by pheromone, cells were collected from mid-log cultures that were not treated with alpha-factor. Cell pellets were snap-frozen in liquid nitrogen and stored at -20 until needed. We extracted the proteins according to previously published methods (45), but scaled up to accommodate the higher cell density. Proteins were resolved by SDS-PAGE on transferred onto nitrocellulose (0.45um, Protran) for Ponceau staining and probing with mouse anti-HA antibodies (1:4000 dilution, Sigma Aldrich) followed by goat anti-mouse HRP-coupled IgG (1:30,000 diluation, Thermo Fisher Scientific). Ponceau S stain of transferred proteins was used for sample normalization and Westerns were visualized using a chemiluminescent HRP antibody detection reagent (Amersham). ImageJ was used for all quantification of bands. Replicate images were acquired and results were averaged across replicates.

### Protein decay

We performed five biological replicates comparing the protein decay rates between the *FIG1*::yECitrine strains YSR0115 (S288c) and YSR0104 (YJM145), the *FUS3*::yECitrine strains YSR0097 (S288c) and YSR0098 (YJM145), the *INA1*::yECitrine strains YSR0143 (S288c) and YSR0144 (YJM145), and the *TOS6*::yECitrine strains YSR0122 (S288c) and YSR0123 (YJM145). We added alpha-factor (50nM final) to mid-log cultures of *FIG1*, *FUS3*, and *TOS6* strains, incubated cultures for three hours, and collected cells from the culture as described above in “Protein sample collection”. For *INA1* strains we collected cells from mid-log cultures as described above in “Protein sample collection”. To halt protein synthesis and observe protein decay, we added cycloheximide (20µg/ml final) to sample wells(46). Twenty minutes after the addition of cycloheximide, we began imaging a new position in the well every ~10 minutes using the approach described above in “Protein imaging and quantification”. We performed two controls using YSR0115, the S288c *FIG1*::yECitrine strain, to verify the alpha-factor and cycloheximide were functioning properly. To mid-log YSR0115 we added alpha-factor and cycloheximide simultaneously, imaged as the samples were imaged, and confirmed cycloheximide function by never observing induction of *FIG1* expression. And to mid-log YSR0115 we added alpha-factor, and as the samples were imaged, and confirmed alpha-factor function by observing normal induction of *FIG1* expression. We estimated the decay rate parameter (lambda) for each gene in each strain by linear regression of the log of the mean over cells of the mean mean pixel intensity across time-points in the cycloheximide treatment. We used a paired t-test, comparing the log ratio of the lambda values between the strains across biological replicates to an expected value of zero. Because *FIG1*, *INA1*, and *TOS6* lambda values were all close to zero and included both positive and negative values, we transformed the *FIG1* lambda values by adding 1 to each.

### Cis-acting vs trans-acting

We measured allele-specific protein expression using fluorescence microscopy with two different fluorophore colors in translational fusions with alleles in the same strain. For each of *FIG1* and *INA1*, starting with the two Citrine strains, we replaced the already deleted ura3 open reading frame with a second copy of *FIG1* or *INA1*. The added DNA included the upstream noncoding region of the gene, the coding sequence of the gene in a c-terminal translational fusion with the blue fluorescent protein gene yECerulean, a 12xMyc tag, and the KanMX G418 resistance gene. We either constructed the cassettes in a plasmid containing 1kb sequences flanking the ura3 open reading frame that was then linearized and transformed in the Citrine strains or we constructed and transformed the cassettes simultaneously using recombineering(47). The latter proved more efficient and robust. We verified integration using PCR, sequencing, and fluorescence microscopy. Using these approaches we constructed eight citrine/cerulean strains: YSR0147 (S288c FIG1S288c::yECitrine FIG1_S288c_::yECerulean), YSR0148 (S288c FIG1_S288c_::yECitrine FIG1_YJM145_::yECerulean), YSR0149 (YJM145 FIG1_YJM145_::yECitrine FIG1_S288c_::yECerulean), YSR0150 (YJM145 FIG1_YJM145_::yECitrine FIG1_YJM145_::yECerulean), YDP0018 (S288c INA1_S288c_::yECitrine INA1_S288c_::yECerulean), YDP0019 (S288c INA1_S288c_::yECitrine INA1_YJM145_::yECerulean), YDP0020 (YJM145 INA1_YJM145_::yECitrine INA1_S288c_::yECerulean), and YDP0021 (YJM145 INA1_YJM145_::yECitrine INA1_YJM145_::yECerulean). For *FIG1* we treated with alpha-factor and performed time-course imaging experiments as described above but now imaging both yECitrine and yECerulean. We performed five biological replicates for *FIG1* (two data points were removed for one of the replicates due to imaging errors), and three biological replicates for *INA1*. For *INA1* we did not treat with alpha-factor and imaged mid-log cultures. We used the strains with different alleles in fusions with yECitrine and yECerulean to measure allele-specific expression and we normalized for fluorophore intensity and genomic location using the strains with the same allele in fusions with yECitrine and yECerulean. For example, we calculated the following for *FIG1*:

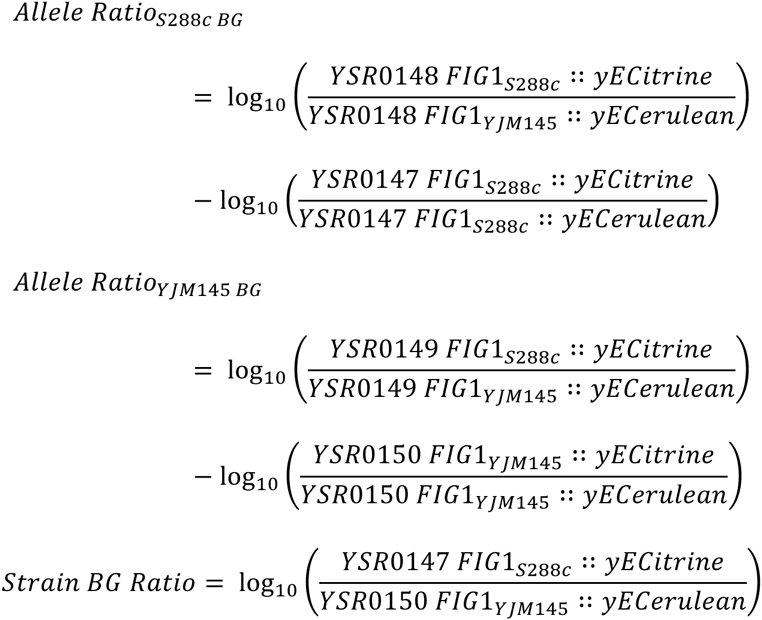

## Acknowledgements

This work was supported by a Human Frontiers Science Program Young Investigator award to SAR (HFSP RGY0073/2010), the San Diego Center for Systems Biology (NIH P50 GM085764), and a National Science Foundation grant to DAP (MCB-1518314) and SAR (MCB-1517482). We thank Molly Burke, Lawrence Du, Sidney Kuo, Isaac Lopez, Ruth Schwartz, Sarah Stockwell, Sharon Tracy, and Allison Wu for technical and conceptual discussions. We thank Katia Bonaldi and Dawn Nagel for input on RT-qPCR. We thank Marketa Ricicova and Carl Hansen for assistance with selecting pheromone concentration and input on cell segmentation. We thank John Houser for input on protein decay assays. We thank Lin Chao, Maitreya Dunham, Mark Estelle, Craig Moyer, Lynn Pillitteri, Lorraine Pillus, José Pruneda-Paz, Clint Spiegel, and Patricia Wittkopp for sharing equipment and reagents.

---

**S1 Figure.**
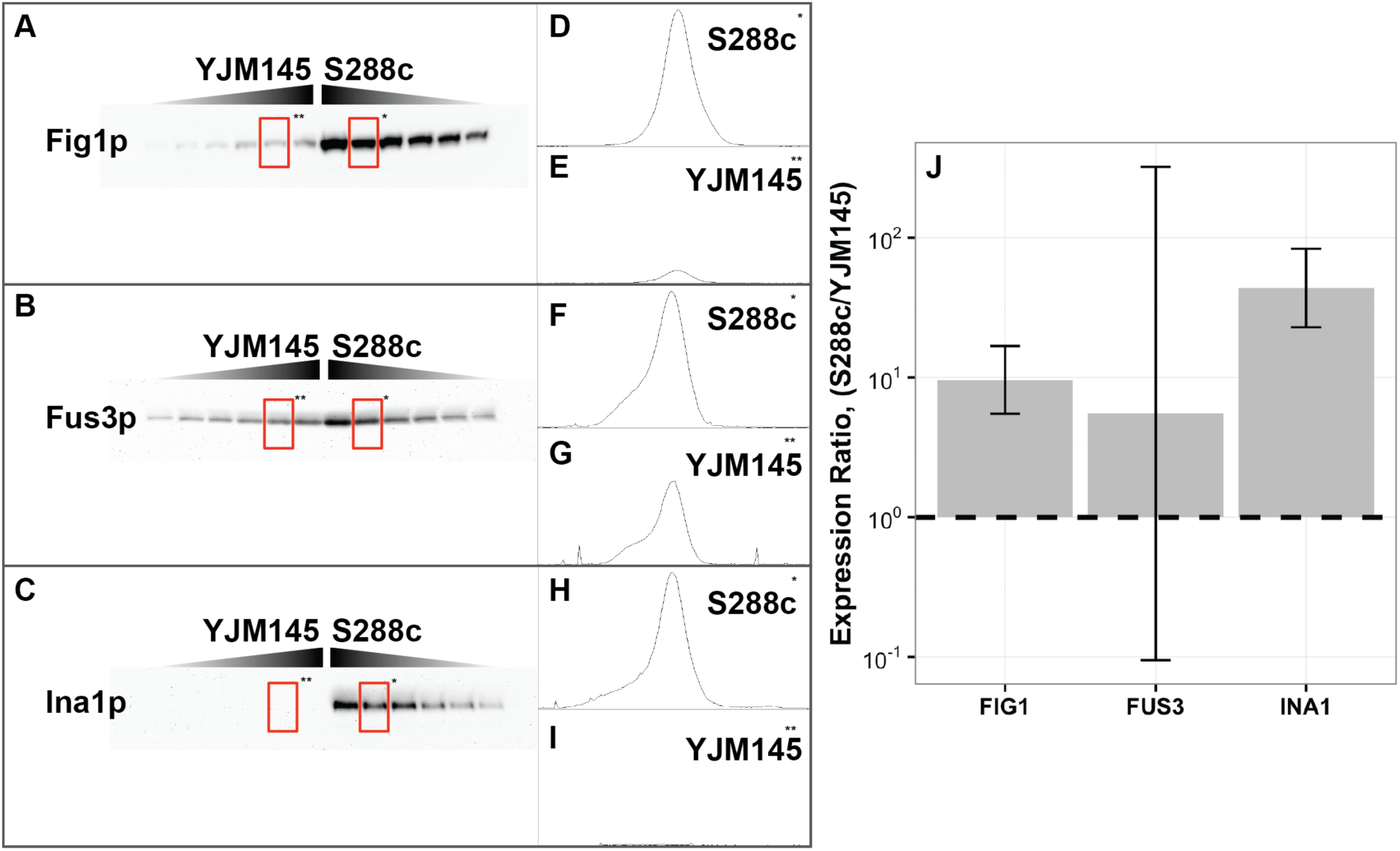
Confirmation of protein expression divergences by Western blot. (A-C) Example western blots for Fig1p (A), Fus3p (B), and Ina1p (C). Two centermost lanes represent an undiluted amount of protein for each strain. Each lane thereafter has been serially diluted by 20%. Lanes marked by an asterisk were used to make the comparisons with the microscopy. (D-I) Band intensity profiles for adjacent boxed blot bands. Pixel intensities were estimated by selecting the area under the peaks using ImageJ’s gel analyzer tool(48). (J) Mean ratio of protein expression between strains over biological replicates. Error bars are 95% confidence intervals.

**Fig S2.**
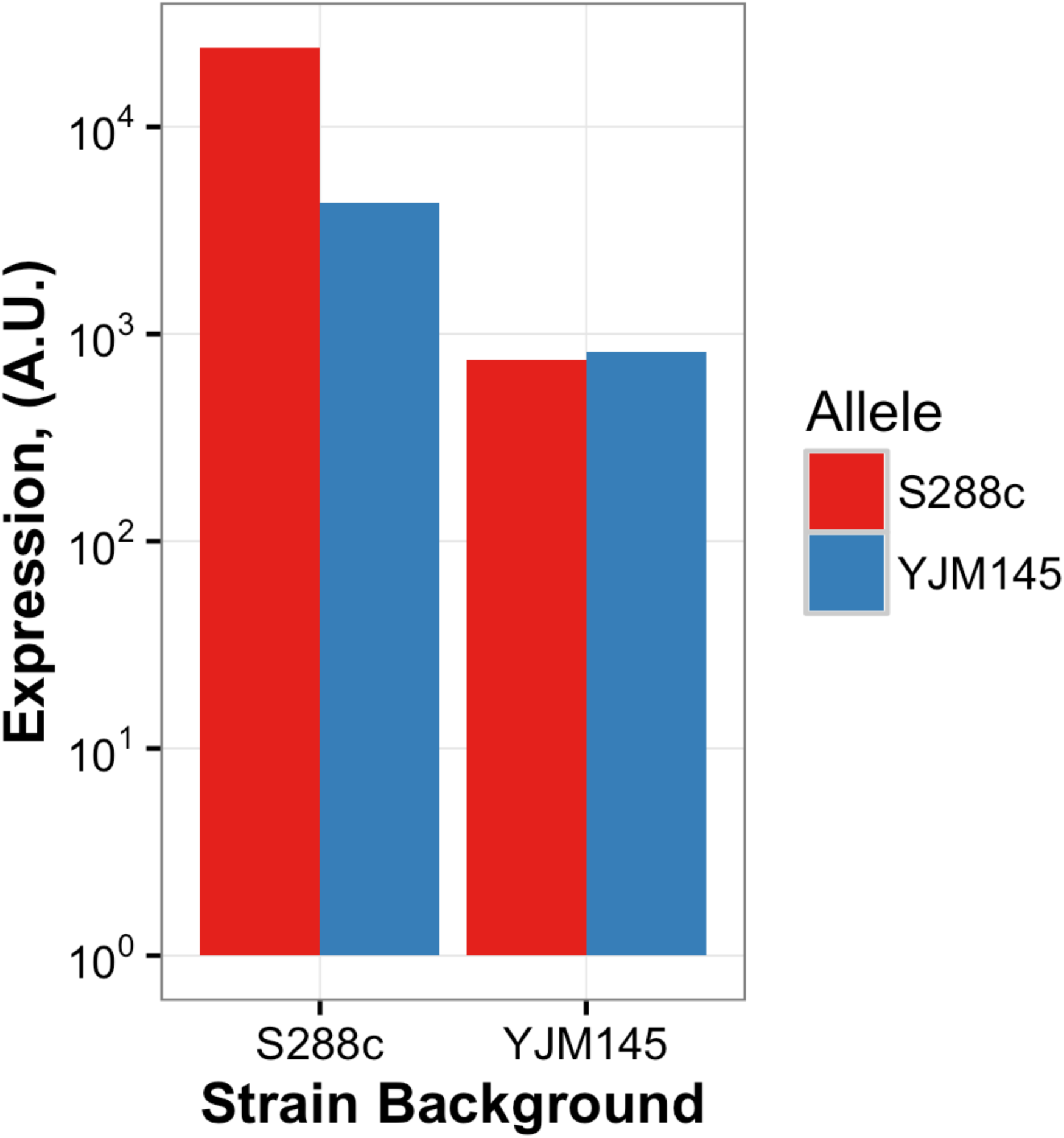
Example allele-specific *INA1* protein expression. Example expression levels of alleles of the *INA1* locus from strains S288c and YJM145 in each of the two strain backgrounds. Y-axis is log10 scale.

